# The mitochondrial transaminase GOT2 supports Japanese encephalitis virus replication in neuronal cells

**DOI:** 10.64898/2026.10.12.758629

**Authors:** Vikas Sood, Anoop Ambikan, Sakshi Khera, Yogita Rawat, Sofja Poznakovs, Teresa Stellaccio, Marieke Boecker, Ákos Végvári, Magda Lourda, Birgitta Lindqvist, Soham Gupta, Iva Filipovic, Manjula Kalia, Ujjwal Neogi

## Abstract

Japanese encephalitis virus (JEV) is the leading cause of viral encephalitis in Asia, yet the host pathways that sustain its replication in the central nervous system remain poorly defined, and no antiviral therapies exist. Here we combine time-resolved transcriptomics and proteomics in mouse neuroblastoma (Neuro-2a) cells with time-resolved proteomics of infected mouse brain, integrated using directionality-penalized p-value merging. JEV infection was accompanied by progressive induction of interferon-stimulated proteins and by changes in the abundance of enzymes of central carbon and energy metabolism, spanning the tricarboxylic acid cycle, glycolysis, oxidative phosphorylation and glutaminolysis. Temporal clustering of the brain proteome linked oxidative phosphorylation remodelling to a dependence on glutamine metabolism. Glutamine antagonism with DON reduced viral RNA, whereas glycolytic inhibition and AKT modulation did not significantly alter it under the conditions tested. Silencing of the mitochondrial transaminase GOT2 reduced JEV RNA, viral protein, and infectious virus production in human SH-SY5Y cells. Our findings identify a glutamine-GOT2 axis as a metabolic dependency of JEV and a candidate host-directed antiviral target.

## Introduction

Japanese Encephalitis Virus (JEV) is a zoonotic, positive-sense single-stranded RNA virus primarily transmitted through the bite of infected mosquitoes. It is a neurotropic flavivirus that causes Japanese encephalitis (JE), a potentially fatal neurological disease. JE is the leading cause of viral encephalitis in Asia, with an estimated 68000 cases reported annually, predominantly affecting children. While most of the cases of JEV infection are asymptomatic, severe infections lead to encephalitis with a high case fatality ratio of 20-30%.^1^ Clinical symptoms of the viral infection include high fever, chills, muscle pain, mental confusion, and convulsions.^2^ Among survivors of Japanese encephalitis, 30–50% develop long-term neurological, cognitive, or psychiatric sequelae, which may result in substantial and potentially lifelong disability.^3^ Currently, there are no antiviral therapies against the virus, and vaccination is the only prophylaxis available.^1^

Following the infected mosquito bite, JEV initially replicates in peripheral tissues and lymphoid organs before entering the bloodstream. Although most infections remain asymptomatic, in a small proportion of individuals the virus invades the central nervous system, where infection of neurons and activation of microglia and astrocytes trigger a strong neuroinflammatory response.^4^ Excessive production of pro-inflammatory cytokines and chemokines can amplify neuronal injury and contribute substantially to disease severity.^5^ However, the molecular mechanisms determining why only a subset of infected individuals develop severe encephalitis remain poorly understood, suggesting that host-intrinsic genetic, immunological, and metabolic factors may critically influence susceptibility to neuroinvasion and neurological damage.

As obligate parasites, viruses reprogram the host’s transcriptional, translational, and metabolic machinery to enable productive viral replication. Accumulating evidence from diverse viral infections demonstrates that immune activation is tightly coupled to profound metabolic rewiring, particularly involving central carbon metabolism, mitochondrial energetics, and amino acid utilization.^6^ Such immunometabolic adaptations not only fuel the biosynthetic demands of viral replication but also critically shape innate and adaptive immune cell effector functions. In neurotropic viral infections, these metabolic perturbations may be further amplified due to the high energy requirements of neurons and the unique immune environment of the central nervous system.^7^ Nevertheless, the extent to which coordinated immune and metabolic reprogramming governs JEV pathogenesis at a systems level remains unexplored.

Systems biology approaches that integrate multiple layers of molecular information offer a powerful strategy to overcome these limitations.^8^ Multi-omics integration enables the construction of regulatory and functional networks that capture the coordinated behavior of genes, proteins, and pathways in a biologically meaningful manner.^9-11^ Network-based analyses can further identify key molecular hubs, upstream regulators, and bottleneck pathways that orchestrate complex host responses.^10^ In the context of viral infections, such approaches have revealed that disease severity is often governed not by isolated pathways but by tightly interconnected immune-metabolic networks that evolve dynamically over the course of infection. In our previous work, we have successfully integrated multi-omics-based approaches against several medically important viruses, including SARS-CoV-2,^11,12^ Crimean-Congo hemorrhagic fever viruses (CCHFV),^10,13^ and Parechoviruses.^14^ Yet, comprehensive integrative multi-omics studies focusing on JEV are still lacking.

Neurons have limited capacity to switch fuel sources, so a neurotropic virus must either tolerate or reshape the metabolism of the cell it replicates in. We therefore asked which host metabolic node JEV requires in neurons, and whether that requirement holds in both the infected murine brain and human neural cells. Combining time-resolved transcriptomics and proteomics with targeted pharmacological and genetic perturbation, we identify the mitochondrial transaminase GOT2 as a host factor supporting JEV replication.

## Results

### Integrated proteomic and transcriptomic analysis reveals progressive metabolic remodelling during infection

To understand the host immune-pathogenic response to JEV infection, mouse Neuro2a (N2a) cells were infected with JEV at an MOI of 5 and harvested at 12, 24, and 48 hours post-infection (hpi) for integrated transcriptomics and LC-MS-based proteomics analysis (Fig. 1A). The viral infection was confirmed by qPCR (Fig 1B) and confocal microscopy (Fig. 1C). Transcriptomic and proteomics datasets were first analysed independently to identify differentially expressed features at each time point relative to mock-infected controls. To capture molecular changes supported across both data layers, the datasets were subsequently integrated using directionality-penalized p-value merging (DPM).^15^ This approach combines differential-expression p-values from the proteomic and transcriptomic analyses while penalizing features for which the direction of change is discordant between the two modalities. The integrated analysis identified 4,973, 7,619 and 6,992 significant genes (p<0.05) at 12, 24 and 48 hpi, respectively (Fig. 1D and Supplementary Table S1). For comparison, p-values were also integrated using Brown’s method, which does not account for the direction of expression changes. To define the biological processes associated with the late-stage response, the 6,992 genes identified by DPM at 48 hpi (p<0.05) were subjected to pathway enrichment analysis. This identified multiple significantly enriched pathways (adjusted p<0.05; Fig. 1E and Supplementary Table S2), predominantly associated with metabolism, signal transduction, and organismal systems. Central carbon metabolic pathways, including the tricarboxylic acid (TCA) cycle, pentose phosphate pathway, pyruvate metabolism and glycolysis, were among the most strongly enriched on the basis of statistical significance and gene overlap, indicating extensive metabolic remodelling at 48hpi. We next examined the abundance profiles of significantly altered proteins associated with the enriched metabolic pathways (adjusted p<0.05). At 12hpi, protein abundance patterns were broadly similar between infected and mock-infected samples. By contrast, increasing separation between the two conditions was apparent at 24 hpi and became more pronounced at 48hpi (Fig. 1F and Supplementary Table S3). Proteins associated with the TCA cycle, pyruvate metabolism, oxidative phosphorylation, and alanine, aspartate, and glutamate metabolism showed particularly prominent increases in abundance at 48hpi. Together, these results indicate a time-dependent increase in the abundance of enzymes in central carbon and energy-producing pathways during the later stages of infection. Mapping of the enriched metabolic pathways onto a simplified pathway framework highlighted changes across glycolysis, the TCA cycle, oxidative phosphorylation, and glutaminolysis (Fig. 1G). Several enzymes connecting these pathways, including GOT1, GOT2, GLS and GLUD1, were increased in abundance (state direction and fold change for each), indicating altered levels of enzymes in central carbon and amino-acid metabolism. The proteomic response was also accompanied by a time-dependent increase in interferon-responsive proteins. At 24hpi, relatively few ISG-associated proteins showed marked changes, whereas by 48 hpi, numerous antiviral proteins, including IFIT1, ISG15, IFITM3 and STAT1, were significantly increased in infected cells (Fig. 1H and Supplementary Tables S4 and S5). Thus, the late infection response was characterized by the simultaneous emergence of pronounced metabolic and antiviral proteomic signatures. Together, these findings reveal a coordinated, time-dependent host response to JEV infection, characterized by extensive metabolic remodeling alongside the progressive induction of antiviral defense programs.

**Figure 1:**
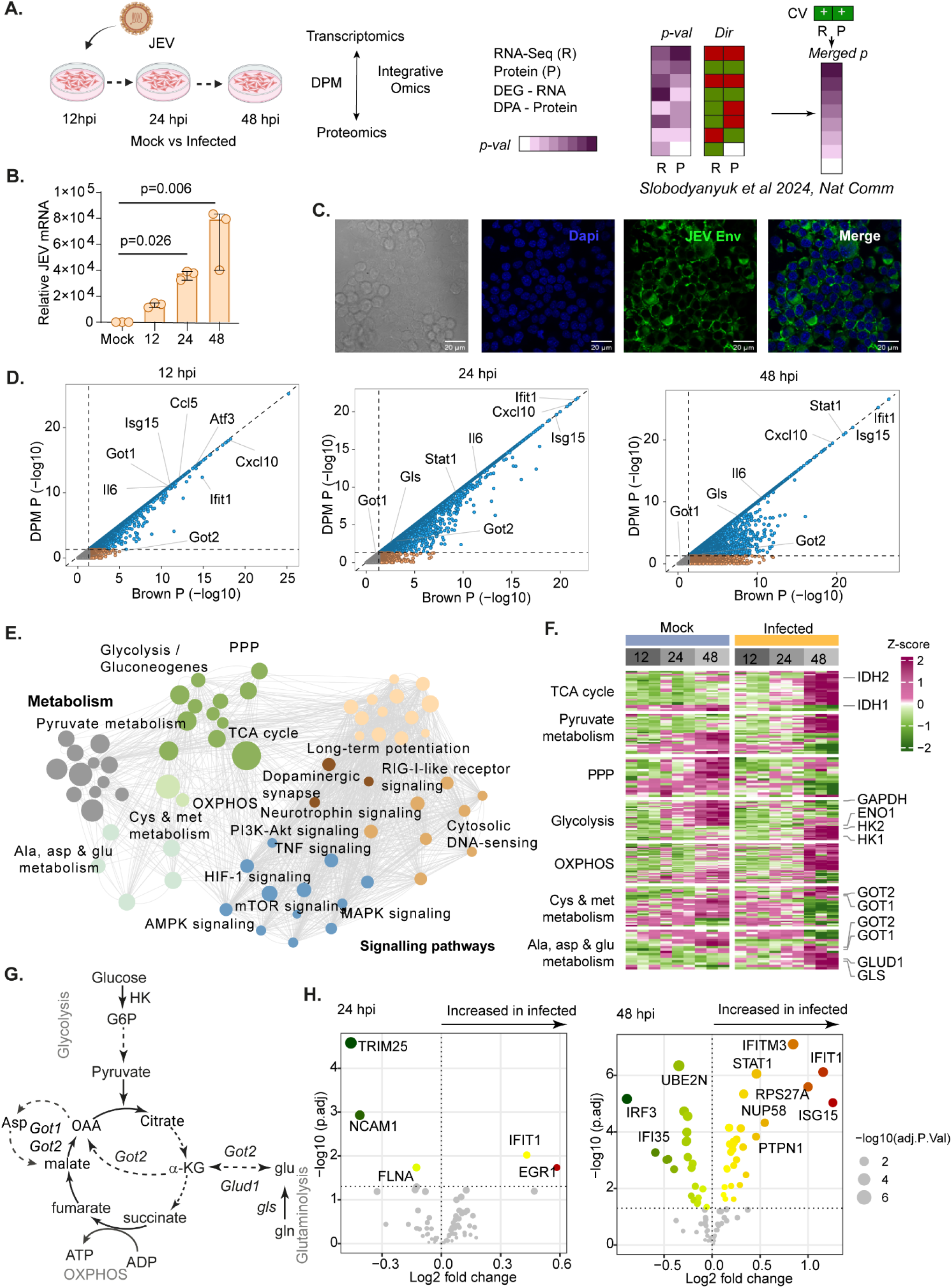
**A)** Experimental design. Mouse Neuroblastome cells (N2a) were infected with JEV at 5MOI harvested at different time points and the levels of JEV were checked using B) Relative JEV mRNA levels C) Representative confocal microscopy images showing JEV envelope protein expression detected using the 4G2 antibody conjugated with Alexa Fluor 488 (green) and nuclei were counterstained with DAPI (blue);Scale bars, 50 µm. D) Scatter plot showing directionality-penalized merged expression difference p-values (DPM p) alongside non-directional merged p-values (Brown’s method), derived from directional integration of proteomic and transcriptomic differential analyses at 12, 24, and 48 hpi. Key genes are labeled. Non-significant genes (p ≥ 0.05) are shown in grey. **E**) Network diagram depicting pathway enrichment results at 48 hpi following directional integration of proteomic and transcriptomic data. The network displays significantly enriched pathways (adjusted p < 0.05). Each node represents a pathway; edges indicate gene overlap between pathways; node size reflects the proportion of significant genes within each pathway. **F**) Heatmap illustrating protein abundance profiles for genes associated with key enriched pathways across all samples, based on proteomics data. Columns denote samples; rows represent proteins, and column annotations indicate sample groups and post-infection time points. **G**) Schematic representation of metabolic pathways highlighted by the integrated multi-omics analysis, illustrating connections between glycolysis, the TCA cycle, oxidative phosphorylation (OXPHOS) and glutaminolysis. Selected enzymes supported by the omics data are indicated at their corresponding metabolic steps. **H**) Volcano plots showing differential abundance of interferon-stimulated genes (ISGs) at 24 and 48 hpi from proteomics data. ISGs were subset from the whole differential proteomics dataset. Non-significant proteins (adjusted p ≥ 0.05) are shown in grey.

### Mouse brain proteomics reveals progressive antiviral activation and metabolic reprogramming during JEV infection

To define the temporal molecular response to JEV infection in vivo, brain tissues collected from infected and mock-treated mice between 1 and 6 days post-infection (dpi) were subjected to proteomic analysis. JEV RNA levels in the brain increased over the course of infection, with a marked rise from 3 dpi onwards and sustained viral burden through 6 dpi (Fig. 2A). Consistent with these kinetics, proteomic profiling revealed limited differences between infected and mock brains during the early stages of infection, followed by increasingly pronounced changes at later time points (Fig. 2B, and Supplementary Table S6). Proteins associated with antiviral immunity, glial activation, and cellular metabolism, including ISG15, IFIT3, CXCL10, GFAP, APOA1, and GOT2, showed prominent temporal changes, with many exhibiting their strongest alterations during late infection. Differential abundance analysis at 6 dpi demonstrated a broad proteomic response to infection, characterized predominantly by increased abundance of host-response proteins (Fig. 2C and Supplementary Table S6). Among the most strongly altered proteins were the interferon-stimulated proteins ISG15 and IFIT3, the chemokine CXCL10, and the astrocytic marker GFAP, consistent with robust activation of antiviral and neuroinflammatory responses. Metabolism-associated proteins, including GOT2 and APOA1, were also significantly altered, suggesting that the late-stage host response extended beyond antiviral defense to involve substantial reorganization of metabolic processes. Selected proteomic changes were independently examined by immunoblotting across the infection time course (Fig. 2D). Accumulation of the viral protein JEV-NS1 became evident at later time points and was accompanied by increased abundance of GOT2 and ISG15, particularly from 4 to 6 dpi. These temporal patterns were consistent with the proteomic data and further linked increasing viral burden with the emergence of metabolic and antiviral host responses in the brain. To spatially resolve virus-associated cellular responses in infected brain tissue, sections were examined by super-resolution TIRF microscopy. JEV antigen was detected in close spatial association with MAP2-positive neuronal structures and in regions containing IBA1-positive microglia (Fig. 2E). Higher-magnification views revealed heterogeneous patterns of viral antigen distribution, including JEV-positive neuronal profiles surrounded by or adjacent to IBA1-positive microglial structures, consistent with a localized microglial response around infected cells. Given the proteomic evidence for altered mitochondrial and amino-acid metabolism, we next examined the spatial relationship between JEV antigen and GOT2. Super-resolution imaging showed prominent GOT2 signal in JEV-positive cells, with colocalization between viral antigen and GOT2 quantified (Fig.2F). These observations are consistent with the proteomic data and show that GOT2 is detectable in JEV-positive cells in the infected brain.

**Figure 2:**
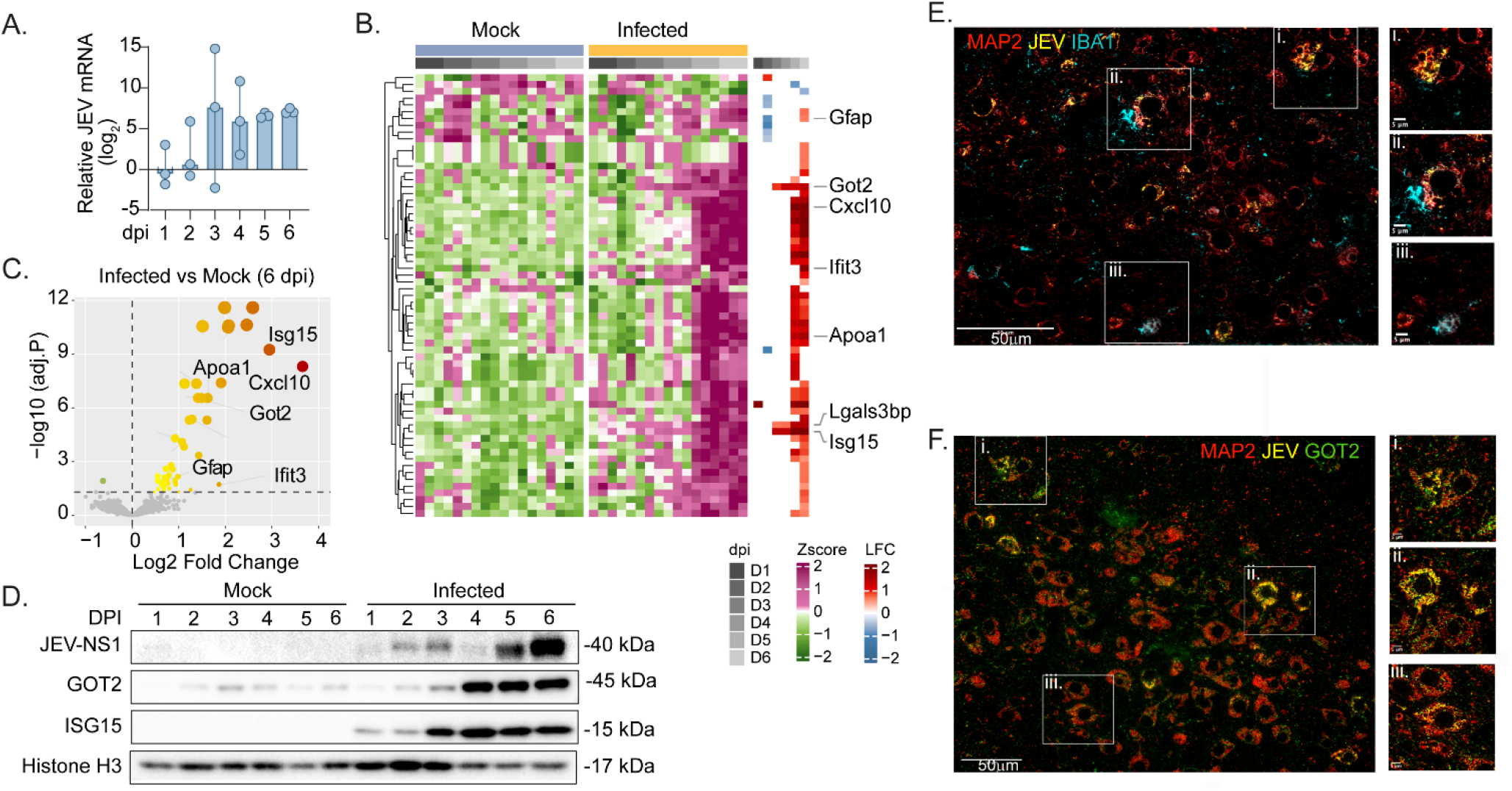
**A)** Viral RNA quantification in the brain. **B)** Heatmap showing the abundance profiles of proteins significantly altered in brain proteomics data (limma, adjusted p < 0.05) in mock versus infected samples at any of the six time points. Column annotations represent the experimental group and time point. Rows are proteins and are annotated with log2 fold-change values from differential abundance analysis at each time point. Cells corresponding to non-significant changes are shown as empty. Selected biologically relevant proteins are labeled. **C**) Volcano plots of differential protein abundance at 6 dpi compared with mock samples. Key proteins are labeled. Non-significant proteins (adjusted p ≥ 0.05) are shown in grey. **D**) Western blot analysis of JEV-NS1, GOT2 and ISG15 along with the HistineH3 as loading control. **E**) Representative super-resolution TIRF microscopy image of infected brain tissue stained for MAP2 (red), JEV antigen (yellow) and IBA1 (cyan). Boxed regions i–iii are shown at higher magnification on the right. Scale bars, 50 µm in the overview image and 5 µm in magnified regions. **F.** Representative super-resolution TIRF microscopy image of infected brain tissue stained for MAP2 (red), JEV antigen (yellow) and GOT2 (green). Boxed regions i–iii are shown at higher magnification on the right. Scale bars, 50 µm in the overview image and 5 µm in magnified regions.

### Temporal proteomic clusters in the infected brain implicate energy metabolism, and glutamine antagonism limits JEV replication

Having defined the spatial context of JEV-associated neuroinflammatory and metabolic alterations, we next examined their temporal organization across the course of infection. To resolve distinct temporal response patterns, we used time series clustering using TMixClust^16^ to identify the trajectories of the protein abundance alterations.^10^ This analysis identified three major clusters with distinct protein dynamics (Fig. 3A and Supplementary Table S7). Cluster 1 showed comparatively modest changes across infection, whereas cluster 2 exhibited a progressive increase in abundance, particularly from 3 dpi onwards. In contrast, cluster 3 displayed a transient increase at 2 dpi followed by a sustained decrease during later infection. These profiles indicate that the brain proteome undergoes temporally structured remodelling rather than a uniform response to increasing viral burden. Pathway enrichment analysis of the temporal clusters revealed distinct biological processes associated with each response trajectory (Fig 3B). Enriched pathways included lipid and cholesterol metabolism, complement and coagulation cascades, leukocyte transendothelial migration, glycolysis and gluconeogenesis, pyruvate metabolism, and oxidative phosphorylation. The co-enrichment of immune-associated and metabolic pathways suggested that progressive neuroinflammatory activation was accompanied by extensive reorganization of energy metabolism during infection. Further, the proteins in glycolysis and oxidative phosphorylation revealed pathway- and time-dependent changes across the course of infection (Fig. 3C). Multiple glycolytic enzymes and components of mitochondrial respiratory-chain complexes I, III and IV were differentially abundant, with distinct patterns emerging across successive stages of infection. Together with the pathway-level analysis, these data indicate coordinated remodelling of cytosolic and mitochondrial energy metabolism in the infected brain. These findings define a temporally coordinated brain response to JEV infection in which increasing viral burden is accompanied by progressive antiviral and neuroinflammatory activation alongside extensive reprogramming of central energy metabolism. To interrogate the functional relevance of glycolysis and OXPHOS during viral infection, we employed a panel of well-characterized metabolic modulators that target distinct nodes of host cellular metabolism. Metabolic inhibitors such as 2-deoxy-D-glucose (2-DG) and 6-diazo-5-oxo-L-norleucine (DON) perturb cellular energy metabolism by interfering with key metabolic pathways that either directly feed into or support the OXPHOS machinery (Fig 3D). Additionally, we used the AKT inhibitor MK-2206 and the AKT activator SC79 to bidirectionally modulate AKT signalling and determine how AKT-dependent regulation of glycolysis and OXPHOS influences cellular susceptibility or resistance to JEV infection. AKT is a central regulator of cellular metabolism that promotes glucose uptake and glycolytic flux while also modulating mitochondrial respiration and OXPHOS.^17,18^ We have used non-toxic concentrations of the drugs (supplementary Fig. S1) to assess their impact on JEV replication. Pharmacological perturbation of the identified metabolic and signalling pathways showed that glutamine antagonism significantly reduced JEV RNA abundance, whereas the other treatments did not produce statistically significant changes under the conditions tested (Fig. 3E). Together, these findings show that glutamine antagonism reduces JEV RNA in neuronal cells, implicating glutamine-dependent metabolism in viral replication.

**Figure 3.**
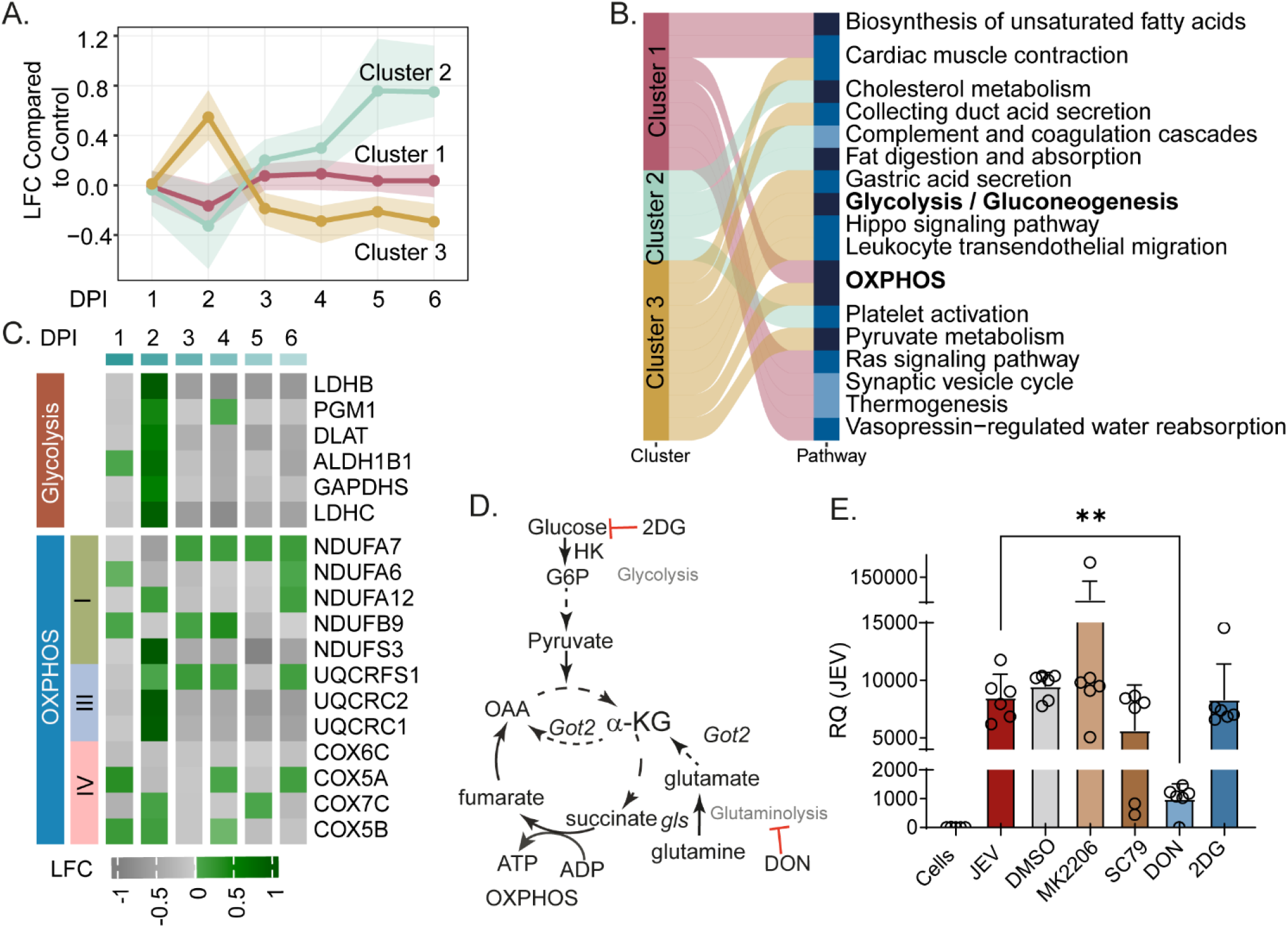
Temporal proteomic profiling reveals coordinated OXPHOS remodelling and a functional requirement for glutamine metabolism during JEV infection. **A.** Temporal expression dynamics clusters identified from the brain proteomics dataset. The line plot displays log2 fold change (LFC) differences in protein abundance at each time point compared to mock. Each line represents a cluster of proteins sharing a characteristic temporal expression pattern. Solid lines indicate the median LFC for each cluster, while shaded regions denote the interquartile range (Q1-Q3) across time points. **B**. Sankey plot illustrating significantly enriched pathways (adjusted p<0.05) associated with the temporal expression clusters. **C**. Heatmap showing significantly abundant proteins (adjusted p < 0.05) belonging to OXPHOS and glycolysis pathways across all time points. Columns represent log2 fold changes in protein abundance at each time point, and row annotations indicate the OXPHOS complex to which each protein belongs. **D**. Schamatic representation of the glycolysis and OXPHOS recation and metabolic inhibitors. **E**. Effect of pharmacological modulation of host metabolic and AKT signalling pathways on JEV replication. JEV-infected cells were treated with DMSO vehicle control, the AKT inhibitor MK-2206, the AKT activator SC79, the glutamine antagonist DON or the glycolysis inhibitor 2-DG. Viral replication was assessed by quantification of relative JEV RNA abundance. Bars show the central tendency and individual points represent biological replicates. DON treatment significantly reduced JEV RNA abundance compared with the DMSO control (**P<0.01), whereas the other treatments did not produce statistically significant changes under the conditions tested.

### A glutamine–glutamate oxaloacetate transaminase 2 (GOT2) metabolic axis supports JEV replication

Pharmacological inhibition with DON implicated glutamine metabolism in supporting JEV replication but did not resolve the specific metabolic node responsible. Given the proteomic alteration of GOT2 and its capacity to link glutamate utilization with TCA-cycle metabolism and mitochondrial respiration, we next performed GOT2 knockdown to test whether this enzyme contributes to the metabolic dependence revealed by DON treatment. To this end, we utilized highly specific small interfering RNA (siRNA) targeting GOT2 to selectively modulate glutamate signaling. siRNA-mediated depletion significantly reduced GOT2 mRNA expression compared with control siRNA-treated cells (p<0.001; Fig. 4A). GOT2 depletion was accompanied by a marked reduction in JEV RNA abundance (p<0.001; Fig. 4B), indicating that GOT2 contributes to efficient viral replication. Consistent with the transcript-level measurements, immunoblot analysis showed reduced GOT2 protein abundance following knockdown, together with a significant decrease in JEV capsid protein levels (p=0.002 and p=0.009, respectively; Fig. 4C). The effect of GOT2 depletion on production of infectious virus was further assessed by plaque assay. Compared with control siRNA-treated cells, GOT2-depleted cells produced substantially fewer plaques, resulting in a significant reduction in plaque-forming units (p<0.001; Fig. 4D). Together, the pharmacological and genetic perturbation data identify a glutamine–GOT2 metabolic axis is required for efficient JEV replication and infectious virus production.

**Figure 4.**
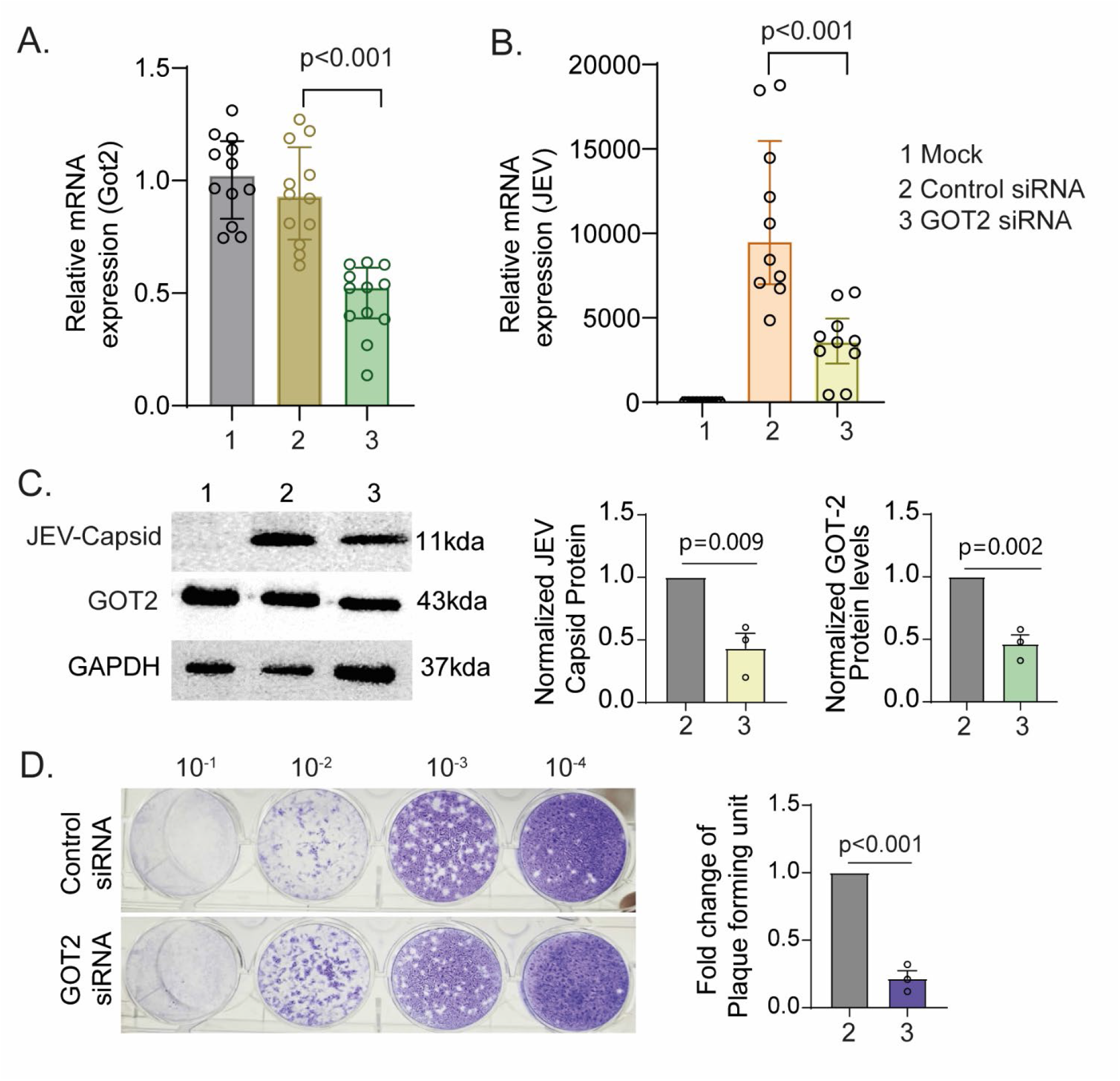
GOT2 depletion suppresses JEV replication and infectious virus production in SH-SY5Y cells at 24hpi (MOI-0.1). **A.** Relative GOT2 mRNA expression in mock, control siRNA and GOT2 siRNA groups. GOT2 knockdown significantly reduced GOT2 transcript abundance compared with control siRNA treatment (p<0.001). Individual points represent biological replicates. **B**. Relative JEV RNA abundance in mock, control siRNA and GOT2 siRNA groups. GOT2 depletion significantly reduced JEV RNA levels compared with control siRNA-treated cells (p<0.001). Individual points represent biological replicates. **C**. Representative immunoblot analysis of JEV capsid and GOT2 protein abundance in mock, control siRNA and GOT2 siRNA groups. GAPDH was used as a loading control. Quantification of JEV capsid and GOT2 protein abundance is shown relative to the control siRNA group. GOT2 depletion significantly reduced both JEV capsid (p=0.009) and GOT2 protein abundance (p=0.002). **D**. Representative plaque assay showing infectious JEV production from control siRNA- and GOT2 siRNA-treated cells across serial dilutions. Quantification of plaque-forming units shows a significant reduction in infectious virus production following GOT2 depletion (p<0.001).

## Discussion

In this study, we integrated multi-omics profiling with functional perturbation experiments to define the temporal host response to JEV infection across both in vitro and in vivo models. Our findings reveal that JEV infection drives a coordinated, time-dependent reprogramming of host cell biology, characterized by the progressive remodeling of central carbon and energy metabolism alongside the induction of antiviral defense programs. Using directionality-penalized integration of proteomic and transcriptomic data, we identified extensive enrichment of TCA cycle, glycolytic, oxidative phosphorylation, and glutaminolysis pathways at late stages of infection, a pattern recapitulated in vivo through temporal proteomic profiling of JEV-infected mouse brain tissue. Critically, functional interrogation of these metabolic alterations identified glutaminolysis as a dependency that actively supports viral replication, with pharmacological inhibition by DON and siRNA-mediated depletion of the mitochondrial transaminase GOT2 both resulting in significantly reduced JEV RNA levels and infectious virus production. These data establish GOT2 as a host factor required for efficient JEV replication in human neural cells and identify glutamine-dependent metabolism as a candidate target for host-directed intervention.

The dominant immunometabolic process during the progressive JEV replication emerging from the integrated multi-omics analysis was extensive remodeling of central carbon and energy metabolism, with enrichment of the TCA cycle, glycolysis, pyruvate metabolism, OXPHOS, and glutaminolysis pathways becoming progressively more pronounced across the infection time course. This metabolic signature was not apparent at early time points but emerged at later time points, suggesting that it reflects active and sustained exploitation of host metabolic resources rather than an acute, non-specific cellular stress response. Metabolic reprogramming during viral infection is increasingly recognized as a conserved strategy by which viruses reshape the host intracellular environment to meet the biosynthetic and energetic demands of replication^19,20^. A recent study further supports this, where metabolomic and multi-omics analyses of JEV-infected mouse brains and neurons revealed enhanced central carbon metabolism, particularly nucleotide synthesis, with upregulation of de novo purine biosynthesis, one-carbon metabolism, and the pentose phosphate pathway.^21^ Similarly, a recent study in Neuro2A cells reported the metabolic landscape of JEV infection, in which increased glycolytic flux and catabolic lipid remodeling were identified as host adaptations supporting viral biosynthetic demands.^22^ While that work established glucose and lipid metabolism as key JEV dependencies, it also described impairment of OXPHOS in the context of glucose utilization, raising the question of how JEV-infected cells sustain mitochondrial TCA cycle activity. Our identification of glutaminolysis as an active viral dependency, and the GOT2-mediated transamination axis as a mechanistic node, provides a potential resolution: glutamine-derived carbon may serve as an anaplerotic substitute, fuelling TCA cycle intermediates independently of glucose oxidation. Taken together, these two complementary studies delineate a multi-arm metabolic reprogramming programme in JEV infection, encompassing aerobic glycolysis, lipid catabolism, and glutaminolysis, each representing a potentially targetable vulnerability in the virus–host interface.

Furthermore, a recent study in Neuro2a cells reported the inhibition of JEV replication following 2-DG treatment. ^23^ No such effect was observed in the human neuroblastoma cell line (SH-SY-5Y) in our study. The divergent antiviral effect of 2-DG observed across model systems is mechanistically informative. While glycolytic inhibition with 2-DG suppressed JEV replication in mouse Neuro-2a cells in a HIF-1α-dependent manner, we have earlier shown that targeting the metabolic network to control viral replication is context-dependent and cell type-specific.^24^ We did not observe a significant effect of 2-DG in SH-SY5Y cells, in contrast to a previous report in mouse Neuro-2a cells. The two experiments differ in species, cell line, MOI, and time point, so this discrepancy cannot be attributed to any one of these factors without a side-by-side comparison. The HIF-1α regulatory axis identified in murine models may orchestrate a broader metabolic reprogramming in human cells that encompasses both glycolytic and glutaminolytic arms, a hypothesis warranting direct experimental interrogation.

The metabolic reprogramming observed across our JEV studies reflects a conserved strategy within the *Flaviviridae* family. Dengue virus (DENV) has been shown to strikingly alter central carbon metabolism, particularly glycolysis, during the course of infection, with increased glucose consumption driving biosynthetic and energetic demands for viral replication, and inhibition of glycolysis significantly blocks infectious DENV production^25^. Zika virus (ZIKV) similarly promotes increased glucose utilisation and glycolytic flux in host cells, while glycolytic rewiring simultaneously results in impaired OXPHOS and mitochondrial dysfunction, particularly in pathogenic ZIKV strains ^26,27^, a pattern strikingly consistent with the OXPHOS impairment and TCA cycle enrichment we identify in JEV-infected cells. Crucially, when glycolysis is pharmacologically blocked during ZIKV infection, inhibition of glutaminolysis prevents the transient recovery of viral protein levels and infectious progeny production, indicating that carbon sources beyond glucose, including glutamine, are required to fuel viral replication^28^, directly paralleling our identification of the glutamine–GOT2 axis as a JEV replication dependency. For the neurotropic West Nile virus (WNV), viral RNA replication is dependent on fatty acid synthesis, as demonstrated by the inhibitory effects of pharmacological blockade of fatty acid synthase^29^. Collectively, these observations across DENV, ZIKV, WNV, and JEV suggest that metabolic flexibility, encompassing coordinated upregulation of glycolysis, glutaminolysis, and lipid catabolism, represents a pan-flavivirus replication strategy, with the relative dependence on each arm varying by virus strain, host species, and cell type, and positioning broad-spectrum metabolic inhibitors for cross-flavivirus antiviral development.

While our study provides a comprehensive multi-omics framework for understanding JEV-induced metabolic reprogramming in mouse cell lines, several limitations warrant consideration. The absence of direct metabolite measurements means that flux through glutaminolysis, glycolysis, and the TCA cycle is inferred from proteomic and transcriptomic enrichment rather than directly quantified. The in vitro findings were derived from human neural cell models rather than primary neurons, and the in vivo temporal proteomic profiling of mouse brain tissue captures a heterogeneous mixture of cell types, precluding cell-type-specific attribution of the observed metabolic reprogramming without single-cell resolution approaches. The species discordance between human in vitro and murine in vivo models is further underscored by the divergent antiviral response to 2-DG observed across model systems, and while we propose glutaminolysis as a compensatory mechanism in human cells, a direct experimental test of this hypothesis through combined glycolytic and glutaminolytic blockade was not performed in the current study. Additionally, although both DON treatment and siRNA-mediated GOT2 depletion reduced viral replication, DON’s broad glutamine antagonism limits mechanistic specificity, and the precise molecular basis by which GOT2-mediated transamination supports JEV replication, whether energetic, biosynthetic, or structural, remains to be fully resolved.

In conclusion, our study provides evidence that the convergence of findings across human and murine systems is further underscored by the deep evolutionary conservation of GOT2 itself, a pyridoxal phosphate-dependent mitochondrial transaminase for JEV replication. Taken together, the temporal multi-omics architecture and dual-modality functional interrogation presented in this study identify GOT2 as a host factor supporting JEV replication and a candidate target for host-directed antiviral development.

## Methods

### Cell culture

Neuro2a (Mouse neuroblastoma) cell line was procured by ATCC (Cat#CCL-131), while C6/36 (*Aedes albopictus*) and BHK21 (Baby Hamster Kidney fibroblast cells) cells were obtained from cell repository at the National Center for Cell Sciences Pune, India. The Neuro2a and BHK21 cell lines were maintained in Dulbecco’s modified Eagle’s medium (DMEM; Lonza Cat#12-604F) supplemented with 10% fetal bovine serum (FBS; Himedia; Cat#RM1112) and 1% antibiotic antimycotic solution (Himedia; Cat#A002A) at 37 in 5% CO_2_. The L-15 medium (Himedia; Cat#AL011A) containing 10% FBS with 1% antibiotic antimycotic solution was used to culture C6/36 cells at 28°C without CO_2_. A hybridoma cell line (D1-4G2-4-15, ATCC; HB-112) was maintained in hybri-Care medium (cat# ATCC 46-X) for generation of mouse monoclonal antibody 4G2 against pan-flavivirus envelope protein.

### Virus generation and quantification

The P20778 strain of Japanese encephalitis virus (JEV) (GenBank accession no. AF080251) was propagated in C6/36 cells. Briefly, the cells were infected with JEV at MOI of 0.1. Then infected cells were incubated at 28°C for three days. The supernatant was collected containing the released virus and it was further stored at -80 °C for long-term storage. For quantification of prepared virus, the BHK-21 cells were seeded into 6 well plate at a density of 0.3x 10^6^ per well and infected with serially diluted JEV virus for 1 hr. Following the incubation, the cells were washed with PBS and 2 ml of DMEM containing 2%FBS, 1% antibiotic, and 1% carboxy methyl cellulose (CMC) were added to each well. The plate was then incubated at 37°C with 5% CO_2_ for 36-48 hours. After incubation, cells were fixed with 10% formaldehyde at room temperature (RT) for 20 minutes. The formaldehyde was removed, and the cells were stained with 0.5% crystal violet solution for 30 minutes. The plate was then rinsed with water to remove excess stain and the plaques formed on the cell monolayer were counted.

### Virus infection in cell line

Neuro2a cells were seeded into the 100 mm dishes at a density of 3×10^6^ cells and incubated overnight. The cells were then infected with JEV at MOI of 5, and uninfected cells were used as control. After 1 hour of incubation at 37°C and 5% CO_2_, cells were washed with PBS and supplemented with fresh DMEM. Cells and supernatant were collected at several time points: 12, 24, and 48 h post-infection.

### Virus infection in mice

Firstly, a mice-adapted strain of JEV S3 was produced in 2-3 days old pups, as previously described.^30^ In the mouse experiment, 3-week-old C57BL/6 mice (n=36) of either sex were weighed and randomly allocated to the infected (n=18) and uninfected groups (n=18). The infected mice were administered with 10^7^ PFU of mouse-adapted JEV intraperitoneally, whereas the uninfected group received an identical volume of incomplete DMEM. The disease progression was evaluated every 24 hours by monitoring for JEV clinical signs, such as weight change, tremors, piloerections, and hind limb paralysis. Three mice from each group (uninfected and infected) were euthanized on each day from the first to the sixth day post infection, and serum, brain, liver, and spleen were collected. For methods: All animal experiments were approved by the Animal Ethics Committee of RCB (RCB/IAEC/2021/099; RCB/IAEC/2024/203). The experiments were conducted in accordance with the Committee for the Purpose of Control and Supervision of Experiments on Animals (CCSEA), Government of India.

### Immunofluorescence microscopy

The cells were seeded on the sterile coverslip precoated with poly-L-lysine and then infected with JEV at MOI of 5. Post-infection, the cells were washed with PBS, fixed with 4% Paraformaldehyde (PFA), and permeabilized with 0.1% triton X-100. Subsequently, the cells were blocked for 1 hour with 1% BSA. Then the cells were incubated with primary anti-flavivirus envelope antibody (in-house) at 1:20 dilution at 4°?C overnight. The cells were washed with PBS followed by staining with Alexa Fluor 488 tagged anti mouse IgG at 1:500 dilution (Cat# A21202) at RT for 1 hr. Lastly, the cells were mounted with Prolong Diamond antifade mountant containing DAPI (Cat# P36966, ThermoFisher). Slides were observed and images were captured on Olympus FV3000 confocal microscope.

### Total internal reflection fluorescence (TIRF) microscopy

Formalin-fixed paraffin-embedded mouse brain tissue was sectioned at 4 µm onto a photonic chips precoated with poly-L-lysine. Chips were baked in an oven at 60°C for 1 hour. Sections were deparaffinilized and rehydrated as follows: xyline I 5 minutes, xyline II 5 minutes, 95% ethanol 3 minutes, 70% ethanol 3 minutes, 50% ethanol 3 minutes, dH_2_O 2 minutes. Sections were then rinsed in PBS. For antigen retrieval, chips were placed into Coplin jars filled with antigen retrieval buffer (2.94 g trisodium citrate, 1000 mL dH_2_O, 0.5 mL Tween20, pH 6) and incubated in a pressure cooker. After the incubation chips were rinsed with dH_2_O. Sections were permeabilized with 0.2% Triton-X-100 for 10 minutes, followed by blocking with blocking buffer (3% BSA in 0.1M PBS) for 1 hour at RT. Sections were then incubated with antibody cocktails at 4°C overnight in a humidified chamber. The first cocktail included primary neuron antibody MAP2 (catalog #17490-1-AP, Proteintech) conjugated with CoraLite Plus 647 (catalog #KFA003, Proteintech) at 1:100 dilution, activated microglia antibody IBA1 (catalog #10904-1-AP, Proteintech) conjugated with CoraLite Plus 555 (catalog #KFA002, Proteintech) at 1:200 dilution, and JEV NS1 antibody (catalog #GTX633820, GneTex) conjugated with CoraLite Plus 488 (catalog #KFA521, Proteintech) at 1:500 dilution. The second cocktail included primary neuron antibody MAP2 conjugated with CoraLite Plus 647 at 1:100 dilution, GOT2 (catalog #14800-1-AP, Proteintech) conjugated with CoraLite Plus 555 (catalog #FA002, Proteintech) at 1:200 dilution, and JEV NS1 antibody conjugated with CoraLite Plus 488 at 1:500 dilution. Antibodies were conjugated according to the manufacturer’s instructions. The following day, sections were rinsed in PB and coverslips were mounted with an aqueous mounting medium. Chips were observed and images were captured using an ACP ZERO microscopy platform (Chip NanoImaging AS).

### Western blot

Cells were lysed in RIPA lysis buffer (Cat#SKU:89900, ThermoFisher) supplemented with a protease inhibitor cocktail (S8820-20TAB, ThermoFisher). The lysate was then incubated for 15 minutes on ice and subjected to three vortex cycles. The lysate was then centrifuged at 13000 rpm for 15 minutes at 4° C and, the supernatant was transferred into a fresh tube. The protein concentration was determined using the BCA assay kit (Cat#23228; ThermoFisher). Subsequently, cell lysates were mixed with 5X SDS buffer (30% glycerol, 5% β-mercaptoethanol, 0.02% bromophenol blue, 10% SDS, 0.25M Tris-HCl ph6.8) and heated to 95°C for 10 minutes. The 30ug protein was separated by SDS-PAGE and transferred to the Nitrocellulose membrane for immunoblotting. The membrane was then blocked with 5% skimmed milk prepared in TBST (Tris-buffered saline solution with Tween 20) at RT for 1 hr. The membrane was incubated with primary antibody overnight at 4°C followed by three washes and a subsequent 1 hour incubation with horseradish peroxide conjugated secondary antibody. The blots were visualized using an ECL (BMU102-EN, ThermoFisher) substrate method and chemiluminescent signals were captured using a ChemiDoc imaging system (BioRad). Antibodies and dilutions used in the western blots are as follows: GOT2, 1:1000 (Proteintech, 14800-1-AP), Histone H3, 1:1000 (Cell signaling, 9715), ISG15, 1:1000 (Santa Cruz Biotechnology, sc-166755), JEV 1:20 (monoclonal antibody, clone 4G2, produced in-house).

### RNA isolation and real-time (qRT-PCR)

RNA was extracted using Trizol reagent (catalog #15596026, ThermoFisher) following the manufacturer’s protocol and then quantified using a nanodrop spectrophotometer. Subsequently, complementary DNA (cDNA) was synthesized using ImProm-II Reverse Transcriptase enzyme (catalog #A3803, Promega). The mRNA expression of JEV was analyzed using SYBR Green DNA-binding fluorescent dye (catalog #RR420A, ThermoFisher) with a Quantstudio 5 RT-PCR detection system. The obtained data was normalized to the expression level of the housekeeping gene GAPDH using the ΔΔCT method, and the gene expression was presented as fold change. The specific primer sequences used for GAPDH and JEV are provided below:

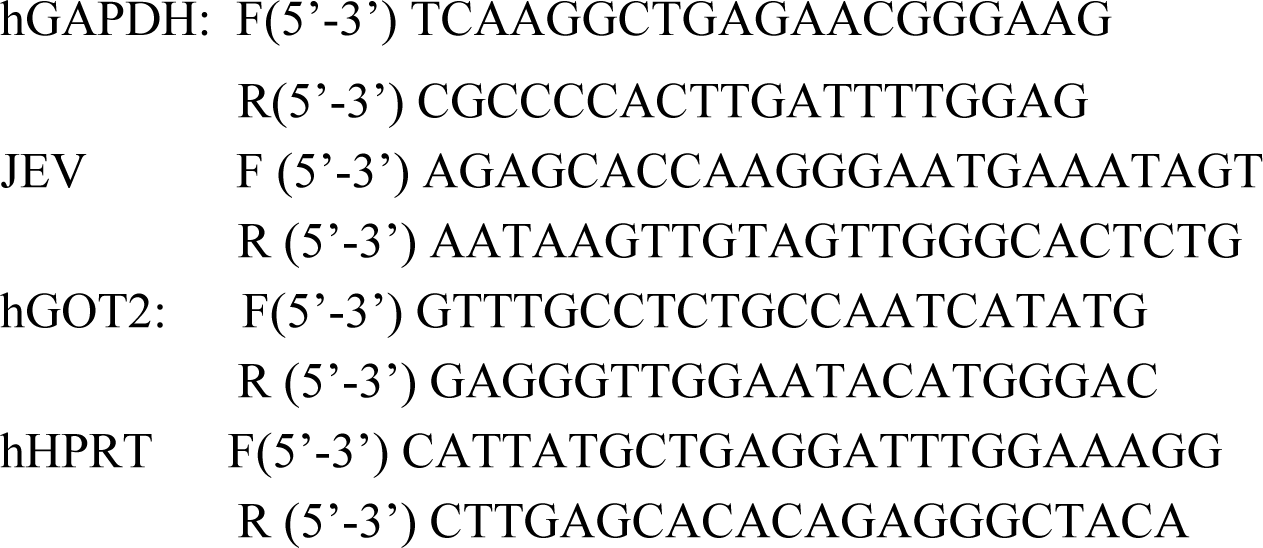

### siRNA transfection

Small interfering RNAs (siRNAs) targeting the *GOT2* gene, along with a non-targeting control siRNA (siNC), were purchased from Dharmacon (ON-TARGETplus). Transfections were carried out using Lipofectamine 3000 (Invitrogen) according to the manufacturer’s protocol. For siRNA transfection, SHSY5Y cells were seeded in 12-well plates and transfected with siRNA at a final concentration of 20 nM, prepared in incomplete media. The cells were then incubated at 37°C to facilitate effective siRNA uptake and gene silencing. Following transfection, the efficiency of gene knockdown was assessed by quantitative PCR (qPCR), western blotting, and plaque assay.

### Plaque assay

The amount of infectious virus released after the siRNA treatment was quantified using a plaque assay, which was performed in the BHK-21 cells. The cells were maintained in T25 in the CO_2_ incubator. On the day of the experiment, 0.1 million cells were seeded in each well of a 12-well plate and allowed to grow for 24 hours. After 24 hours, cells were washed and infected with various doses of the virus ranging from 10^-1^ to 10^-4^. After 1 hour long incubation with the virus, cells were washed and 1ml of DMEM containing 2%FBS, 1% antibiotic, and 1% carboxy methyl cellulose (CMC) was added to each well. The cells were then incubated for 48-96 hours until the plaques became visible. Once the plaques were obtained, 10% formaldehyde solution was added to each well of the 12-well plate for 30 minutes to neutralize the virus particles. Thereafter, the formaldehyde was removed, and the cells were stained with 0.5% crystal violet solution for 30 minutes. The plate was then rinsed with water to remove excess stain and the plaques formed on the cell monolayer were counted manually.

### Transcriptomics

RNA sequencing data were processed using the nf-core/rnaseq workflow (v3.19.0; doi: 10.5281/zenodo.1400710), a community-curated and standardized pipeline from the nf-core initiative.^31^ The workflow was executed using Nextflow v25.04.4,^32^ ensuring scalability and reproducibility of the analysis. All computational environments and software dependencies were managed using Bioconda^33^ and BioContainers,^34^ enabling consistent and reproducible execution across computational platforms. Sequencing reads were aligned to the mouse reference genome GRCm39, obtained from Ensembl. Corresponding gene annotation files were downloaded from Ensembl and used consistently as the reference annotation throughout all RNA-seq analyses. Read alignment and transcript quantification were performed within the nf-core/rnaseq pipeline using the STAR–Salmon mode. In this configuration, STAR,^35^ a splice-aware aligner, was used to map sequencing reads to the reference genome, enabling accurate alignment across exon–exon junctions and complex transcript structures. Following alignment, transcript-level abundance estimation and gene-level quantification were carried out using Salmon in alignment-based mode,^36^ leveraging the STAR-generated alignments. Gene-level expression estimates were obtained by aggregating transcript-level abundances according to the Ensembl gene annotation. For downstream analyses, gene-level estimated counts and TPM values corresponding to Ensembl gene biotypes, specifically protein-coding genes and long non-coding RNAs (lncRNAs) were retained. In total, the analysis included 21,684 protein-coding genes and 11,608 long non-coding RNAs. Gene-level count matrices generated by the pipeline were used for all subsequent analyses. Differential gene expression analysis was performed using the limma R package v3.50.0.^37^ Differential comparisons were conducted between infected and mock samples independently at each time point.

### Proteomics

Proteomic data were initially normalized using the NormalyzerDE R package v1.22.0.^38^ The package evaluates normalization performance using eight different normalization methods and generates validation figures to assist in method selection. Based on inspection of the validation metrics and diagnostic plots, quantile normalization was identified as the most appropriate method and was therefore applied for all downstream analyses. Differential protein abundance analysis was performed using the limma R package v3.50.0.^37^ Differential comparisons were carried out between infected and mock samples at each time point independently.

Brain proteomic data were subjected to missing value imputation and batch-effect correction prior to differential abundance analysis. Missing values were imputed using a k-nearest neighbors (KNN) based approach implemented in the impute R package v1.78.0. Batch effects were corrected using the ComBat method implemented in the sva R package v3.52.0.

Time-series clustering of protein abundance profiles was performed using the TMixClust R package v1.26.0.^39^ Log2 fold-change values derived from the limma analysis, calculated relative to the corresponding mock groups at each time point, were used as input for clustering. The optimal number of clusters was determined based on silhouette width, providing an objective measure of cluster separation and cohesion.

### Proteo-transcriptomics integration

Proteo-transcriptomics integration of the cell line data was performed using Directional P-value Merging (DPM), a method designed for directional integration of multi-omics datasets.^15^ DPM incorporates user-defined directional constraints to prioritize genes or proteins whose directions of change across omics layers are consistent with the specified constraints, while penalizing those with discordant directional signals. The DPM algorithm, implemented in the ActivePathways R package v2.0.6,^40^ available from CRAN, was used for integration. For each time point, statistical significance (p-values) and log2 fold-change values from both transcriptomic and proteomic analyses were provided as input. Brown’s method, which accounts for covariance among correlated tests, was used to merge p-values across datasets.

Gene set enrichment analysis was subsequently performed using the enrichr module implemented in Gseapy v1.1.5^41,42^ to identify significantly enriched biological pathways associated with the integrated multi-omics signals.

### Data visualization

Data visualization was performed using established tools. Scatter plots, volcano plots, and line plots were generated using R package ggplot2 (v3.5.1). Heatmaps were generated using the ComplexHeatmap R package (v2.20.0).^43^ Network visualizations were created using Cytoscape software.

### Statistical analysis

All experiments were performed in three independent biological replicates, unless otherwise stated. Data are presented as mean ± SEM. Statistical analyses were performed using GraphPad Prism (version 8.0.2). For comparisons between two groups, an unpaired two-tailed Student’s t-test was used. For comparisons involving more than two groups, one way ANOVA followed by Tukey’s multiple comparisons post hoc test was employed. A p-value < 0.05 was considered statistically significant.

## Authors’ contributions

Conceptualization:V.S., and U.N.

Conceptualization of the bioinformatics part: A.A., I.F., and U.N.

Methodology: V.S., S.K., A.A., S.P.,T.S., Y.R.,M.B., A.V., M.L., B.L.,

Investigation: M.V., S.B., and N.J.

Visualization: V.S., A.A., and U.N.

Supervision: V.S., M.K., S.G., I.F., and U.N.,

Funding acquisition: V.S., M.K., and U.N.

Writing-original draft: V.S., A.A. and U.N.

Writing-review & editing: All authors

## Data availability

Raw FASTQ files are deposited in NCBI SRA with Bioproject accession number PRJNA1399669. The mass spectrometry proteomics data have been deposited to the ProteomeXchange Consortium via the PRIDE partner repository with the dataset identifier PXD082527. All the codes used for the analysos of proteomics and transcriptomics data are deposited in the github repo: https://github.com/SystemsVirology/JEV_Pathogenesis

## Acknowledgments

UN acknowledges support received from the European Union’s HORIZON Research and Innovation Actions under grant agreement No. 101191315 (COMBAT). Views and opinions expressed are, however, those of the authors only and do not necessarily reflect those of the European Union. The European Union cannot be held responsible for them. UN also acknowledges support received from the Swedish Research Council grant (2021-00993) and Karolinska Institutet Consolidator Grant (2-117/2023). IF acknowledges support received from the Swedish Research Council Starting Grant (2025-02198), Jeanssons Stiftelser (J2025-0113), and Åke Wibergs Stiftelse (M25-0374). VS is thankful for the DBT grant (BT/PR30819/MED/29/1368/2019).

