## Supplementary figures and images for "The mitochondrial transaminase GOT2 supports Japanese encephalitis virus replication in neuronal cells"

### Supplementary Fig S1

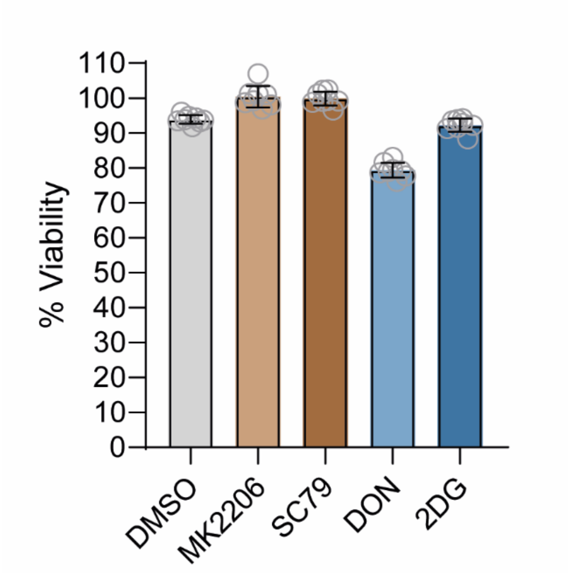


Figure S1. Viability following the drug treatment
